# Phosphorylation of CARM1 at Ser572 preserves muscle stem cell self-renewal by restricting premature myogenic commitment

**DOI:** 10.64898/2026.10.17.759715

**Authors:** Cordell A. VanGenderen, Rebecca Robertson, Jules A. Granet, Ishtiaque Hossain, Ronit Mohapatra, John Saber, Michael A. Rudnicki, William A. Pastor, Natasha C. Chang

## Abstract

Skeletal muscle regeneration depends on satellite cells, which must balance myogenic commitment with self-renewal to maintain regenerative capacity throughout life. The methyltransferase CARM1 is a key regulator of satellite cell fate, acting through methylation of PAX7 to promote *Myf5* expression and myogenic commitment. The dystrophin glycoprotein complex regulates this pathway through p38γ-mediated phosphorylation of CARM1 at Ser572, however, the physiological role of this phosphorylation event *in vivo* remains unknown. Here, we generated a phosphorylation-deficient CARM1 knock-in mouse (CARM1 SA) to determine how CARM1 Ser572 phosphorylation regulates satellite cell fate during regenerative myogenesis. Satellite cell-derived myoblasts from CARM1 SA mice exhibited reduced PAX7 expression, impaired fusion, and abnormal myotube formation despite transcriptional signatures consistent with accelerated myogenic progression. *Ex vivo* single-myofiber assays revealed precocious acquisition of a committed progenitor identity accompanied by a reduction in the self-renewing satellite cell population. While muscle development and regeneration following a single injury were largely preserved, repeated regenerative challenge resulted in progressive depletion of the satellite cell pool in CARM1 SA mice. Transcriptomic analysis demonstrated minimal effects under homeostatic conditions but identified substantial alterations during regeneration, including reduced expression of extracellular matrix-associated genes, impaired TGFβ-SMAD signaling, decreased *Tgfbr3* expression, and increased MYC activity, consistent with a loss of self-renewal programs and enhanced activation. Collectively, these findings demonstrate that phosphorylation of CARM1 at Ser572 is a critical mechanism to delay satellite cell commitment, thereby maintaining a balance between self-renewal and commitment to ensure long-term maintenance of the muscle stem cell compartment.

## Introduction

The regenerative capacity of muscle tissue relies upon a population of resident muscle stem cells, referred to as satellite cells (Lepper et al., 2011; Sambasivan et al., 2011; von Maltzahn et al., 2013). Under homeostatic conditions satellite cells reside in a quiescent state beneath the basal lamina of skeletal muscle tissue, and upon injury become activated to facilitate muscle repair, a process known as regenerative myogenesis (Bentzinger et al., 2012). Following their activation, satellite cells make a fate decision, either committing to the myogenic program and ultimately fusing to repair damaged myofibers or undergoing self-renewal and returning to quiescence to maintain the stem cell pool (Giordani et al., 2018; Millay et al., 2013; Relaix et al., 2021; Zammit et al., 2004). This fate decision is a critical step during regenerative myogenesis, as a balance between both myogenic commitment and self-renewal is necessary to ensure efficient muscle regeneration, while preserving regenerative capacity for future repair (Giordani et al., 2018; Relaix et al., 2021).

Satellite cell fate progression during regenerative myogenesis is mediated by the temporal expression of myogenic transcription factors. The paired box transcription factor PAX7 is expressed in satellite stem cells and regulates transcriptional programs that maintain satellite cell identity (Kuang et al., 2007; McKinnell et al., 2008; Seale et al., 2000). When PAX7 is methylated by the protein methyltransferase CARM1, it recruits the ASH2L:MLL1/2:WDR5:RBBP5 histone H3K4 methyltransferase complex to the promoter region of the myogenic transcription factor *Myf5* (Kawabe et al., 2012). This recruitment promotes H3K4 trimethylation at the *Myf5* promoter to enhance its transcription. This transcriptional activation of *Myf5* by PAX7 signals myogenic commitment of the satellite cells (Kuang et al., 2007; McKinnell et al., 2008). PAX7+/MYF5+ satellite cells will subsequently co-express MYOD and proliferate as committed myogenic progenitors (Cornelison and Wold, 1997). After sufficient proliferation, these progenitors begin to lose PAX7 and gain MYOG expression, a marker of terminal differentiation as they become myocytes, which then fuse with the damaged myofiber to facilitate repair (Zammit, 2017). Alternatively, following proliferation, PAX7+/MYOD+ cells may downregulate MYOD expression and return to quiescence, undergoing self-renewal to maintain the satellite cell pool (Halevy et al., 2004; Zammit et al., 2004). Consistent with self-renewal being an actively regulated fate decision, single-cell chromatin accessibility profiling during muscle regeneration identified a distinct self-renewing satellite cell population characterized by expression of the transforming growth factor β (TGFβ) coreceptor TGFBR3 (betaglycan) (Okafor et al., 2023). In these cells, TGFβ signaling promotes SMAD4-dependent activation of both SMAD2/3 and SMAD1/5 transcriptional programs, supporting satellite cell self-renewal and maintenance of the stem cell pool (Okafor et al., 2023).

Within the satellite cell pool, a subset of cells has never expressed *Myf5*, representing a stem-like population that remains uncommitted while retaining a high capacity for self-renewal (Kuang et al., 2007; Le Grand et al., 2009). Although both PAX7*+/*MYF5*-* and PAX7*+/*MYF5*+* satellite cells are capable of self-renewal, *Myf5* expression demarcates two functionally distinct satellite cell populations: PAX7+/MYF5-satellite cells exhibit superior self-renewal potential and contribute to long-term stem cell maintenance, whereas PAX7+/MYF5+ satellite cells are more committed to the myogenic lineage and display a greater propensity for differentiation (Kuang et al., 2007).

Thus, induction of *Myf5* expression through CARM1-mediated methylation of PAX7 represents a critical step in satellite cell fate determination and must be tightly regulated to maintain the balance between commitment and self-renewal (Chang et al., 2018; Kawabe et al., 2012). CARM1 is expressed as two alternatively spliced isoforms that differ in their C-terminal regions. The shorter isoform, termed CARM1Δ15, lacks exon 15 and is selectively responsible for methylation of PAX7. A key mechanism regulating CARM1-mediated methylation of PAX7 is the dystrophin glycoprotein complex (DGC), which controls the subcellular localization of CARM1Δ15. DGC-associated p38γ MAPK phosphorylates CARM1Δ15 at Ser572, promoting its retention in the cytoplasm and preventing its nuclear interaction with PAX7. Consequently, PAX7 methylation and subsequent *Myf5* induction are suppressed, limiting satellite cell commitment (Chang et al., 2018).

The physiological importance of this regulatory axis is highlighted in murine models of Duchenne muscular dystrophy (DMD). In *mdx* mice, a model of DMD that carries a point mutation in the *Dmd* gene resulting in a premature stop codon and thus loss of dystrophin (Bulfield et al., 1984), p38γ activity is dysregulated resulting in hyperphosphorylation of CARM1. Consequently, PAX7 methylation and satellite cell commitment are impaired, thereby impairing their regenerative capacity (Chang et al., 2018). To directly examine the role of CARM1Δ15 Ser572 phosphorylation in satellite cell fate determination, we generated a phosphorylation-deficient constitutive CARM1 knock-in mouse by CRISPR/Cas9-mediated substitution of Ser572 (CARM1Δ15) / Ser595 (full-length CARM1) with alanine (CARM1 SA). Unlike mice with p38γ deletion or the *mdx* model, this approach isolates the specific contribution of Ser572 phosphorylation without disrupting other p38γ-dependent signaling pathways or introducing the broader pathological changes associated with muscular dystrophy.

CARM1 SA mice and their isolated satellite cells were used to investigate the role of Ser572 phosphorylation in satellite cell fate determination. *In vitro* differentiation of CARM1 SA myogenic progenitors revealed defects in myogenic differentiation. Prior to differentiation, CARM1 SA progenitors exhibited reduced PAX7 expression, indicating a precocious differentiation and premature loss of the stem-like state. Evidence for precocious differentiation was further supported by transcriptional profiling of myogenic progenitors prior to differentiation, which revealed increased expression of genes associated with skeletal muscle development in CARM1 SA mutants. This phenotype was further investigated *ex vivo* using single myofiber culture assays, which demonstrated accelerated acquisition of a committed myoblast identity accompanied by a reduction in the self-renewing stem cell population. Examination of self-renewal *in vivo* using muscle regeneration assays revealed a reduction of the satellite cell pool following repeated injury, indicating impairments to self-renewal. Consistent with this finding, transcriptomic analysis of activated satellite cells three days post-injury demonstrated reduced expression of self-renewal-associated gene programs, including those related to SMAD signaling and extracellular matrix remodeling. Collectively, these findings demonstrate that CARM1 Ser572 phosphorylation delays satellite cell commitment, thereby promoting self-renewal and preserving the balance between self-renewal and myogenic commitment.

## Results

### Generation of CARM1 SA mice

To investigate the functional role of CARM1 phosphorylation at Ser572/595, we generated phosphorylation-deficient CARM1 mutant mice (CARM1 SA). The Ser572/595 residue is encoded by a TCG codon, which was converted to GCC through CRISPR/Cas9-mediated gene editing and homology-directed repair. Specifically, c.T1897/1828T>G and c.G1899/1830G>C substitutions converted the serine codon to alanine in both CARM1 isoforms, thereby preventing phosphorylation at this site (Fig. S1, A and B). Successful genome editing at the *Carm1* locus was confirmed by Sanger sequencing and endpoint PCR (Fig. S1, C and D).

### Methyltransferase activity of CARM1 during in vitro myoblast differentiation

To determine whether inhibition of CARM1 phosphorylation alters its expression or activity during myogenic differentiation, primary myoblasts isolated from wild-type and CARM1 SA mice were differentiated *in vitro* and analyzed for CARM1 abundance, subcellular localization and methyltransferase activity. Myoblasts were cultured to confluence (day 0) and differentiation was induced by serum starvation. In wild-type myoblasts, CARM1 protein abundance was highest under growth conditions (day -2) and progressively declined during differentiation, reaching 30.5% of baseline levels by day 2 (Fig. 1, A and B; p = 0.0166). To assess CARM1 methyltransferase activity, we assessed levels of CARM1-methylated substrates using an α-methyl-NFIB antibody, which recognizes multiple CARM1-methylated substrates (Gao et al., 2023). Consistent with the reduction in CARM1 protein abundance, methylated substrate levels also declined during differentiation, reaching 21.9% of baseline by day four post-differentiation (Fig. 1, A and C; p = 0.0011). Primary myoblasts isolated from CARM1 SA mutant mice exhibited similar temporal declines in both CARM1 abundance and methyltransferase activity, with no overall effect of genotype detected across the differentiation time course (Fig. 1, A-C; p = 0.1356).

**Figure 1.**
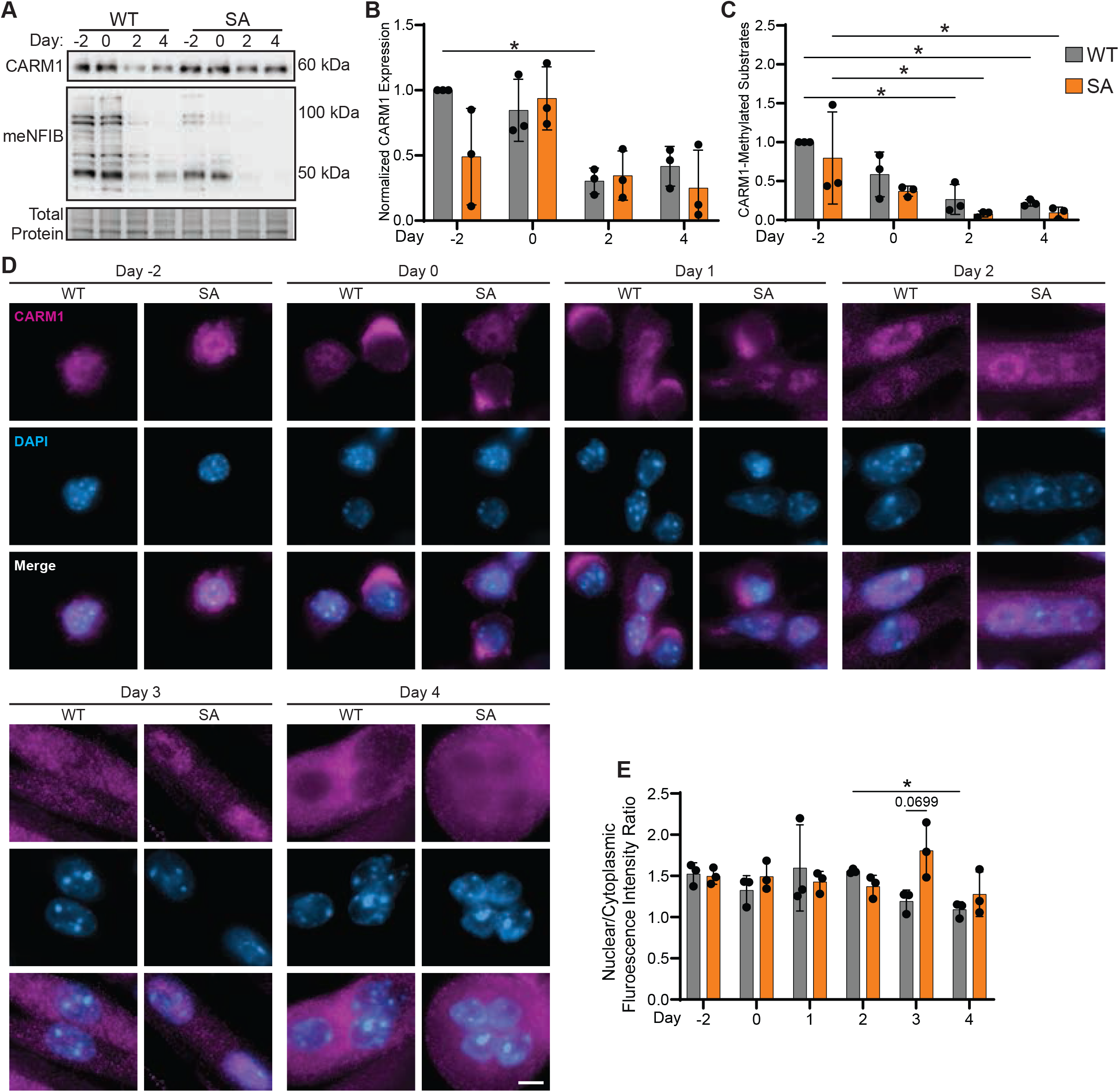
CARM1 expression, localization, and methyltransferase activity during myoblast differentiation. **A)** Primary myoblasts isolated from CARM1 wild-type (WT) and Ser572A (SA) mice were differentiated for four days. CARM1 expression and CARM1-methylated substrates (detected using an anti-methyl-NFIB antibody) were analyzed on days -2, 0, 2, and 4 of differentiation by immunoblot. Quantification of **B)** CARM1 and **C)** CARM1 methylated substrates, normalized to total protein and expressed relative to CARM1 WT at day -2. **D)** CARM1 WT and SA primary myoblasts were differentiated and fixed at the indicated time points. Cells were immunostained for CARM1 (magenta) and nuclei were counterstained with DAPI (blue). Representative images from days -2, 0, 1, 2, 3, and 4 are shown. **E)** CARM1 subcellular localization was quantified as the ratio of nuclear-to-cytoplasmic CARM1 fluorescence intensity. Data are represented as mean ± SD; n = 3 biological replicates; in **D-E** a minimum of 30 cells were analyzed per replicate; *p ≤ 0.05, **p ≤ 0.01; two-way ANOVA with Tukey multiple comparisons. Scale bar represents 5 µm.

Given that phosphorylation regulates CARM1 nuclear localization, we examined CARM1 subcellular localization during myogenic differentiation by immunofluorescent microscopy. Under growth conditions, CARM1 was predominantly localized to the nucleus of wild-type myoblasts (Fig. 1, D and E). During early differentiation (days 0-2), CARM1 remained predominantly nuclear in most cells. However, a subset of undifferentiated myoblasts at days 0 and 1 exhibited pronounced cytoplasmic redistribution of CARM1 (Fig. 1, D and E), suggesting dynamic regulation of CARM1 nucleus to cytoplasmic shuttling during the initiation of differentiation. As differentiation progressed and multinucleated myotubes matured, CARM1 localization shifted increasingly toward the cytoplasm, with the lowest nuclear enrichment observed at day 4 (Fig. 1, D and E). Quantification of the nuclear-to-cytoplasmic fluorescence intensity ratio supported this trend, declining from 1.56 at day 2 to 1.19 at day 3 (p = 0.1383) and becoming significantly reduced to 1.09 at day 4 (Fig. 1 E; p = 0.0465). Although cells exhibiting strong cytoplasmic redistribution were readily observed at days 0 and 1, their frequency was insufficient to significantly alter the average nuclear-to-cytoplasmic fluorescence intensity ratio during early differentiation. Together, these findings demonstrate that CARM1 localization is dynamically regulated throughout myogenic differentiation, undergoing stage-specific nuclear-to-cytoplasmic redistribution that reflects the transition from proliferating myoblasts to mature multinucleated myotubes.

Notably, inhibition of CARM1 phosphorylation in CARM1 SA myoblasts altered this localization pattern, resulting in retention of CARM1 within the nuclei of differentiated myotubes. In contrast to wild-type cells, CARM1 SA myotubes exhibited no significant changes in the nuclear-to-cytoplasmic fluorescence intensity ratio between days 2 and 4 of differentiation. Moreover, at day 3 of differentiation, CARM1 in CARM1 SA myotubes showed a trend toward greater nuclear enrichment compared with wild-type controls (Fig. 1, D and E; p = 0.0699), suggesting a role for CARM1 Ser572 phosphorylation in regulating CARM1 subcellular localization during myotube formation. Collectively, these findings demonstrate that CARM1 abundance and methyltransferase activity are highest in undifferentiated myoblasts and decline during myogenic differentiation. Furthermore, phosphorylation at Ser572 regulates CARM1 nucleocytoplasmic trafficking during differentiation, with loss of this phosphorylation site promoting nuclear retention of CARM1 in differentiated myotubes.

### Inhibition of CARM1 Ser572 phosphorylation impairs myogenic differentiation in vitro

We next examined the impact of inhibiting CARM1 Ser572 phosphorylation on the differentiation of satellite cell-derived myoblasts into multinucleated myotubes *in vitro*. Differentiation was assessed by evaluating myotube morphology and quantifying the fusion index, defined as the percentage of nuclei incorporated into multinucleated myotubes. At two days post-differentiation, the fusion index of CARM1 SA myoblasts was reduced to 77.2% of that observed in wild-type cultures (Fig. 2, A and B; p = 0.0201), indicating impaired myogenic fusion. By four days post-differentiation, CARM1 SA myotubes exhibited marked morphological abnormalities characterized by large, disorganized clusters of myonuclei rather than the elongated myotubes observed in wild-type cultures (Fig. 2 C).

**Figure 2.**
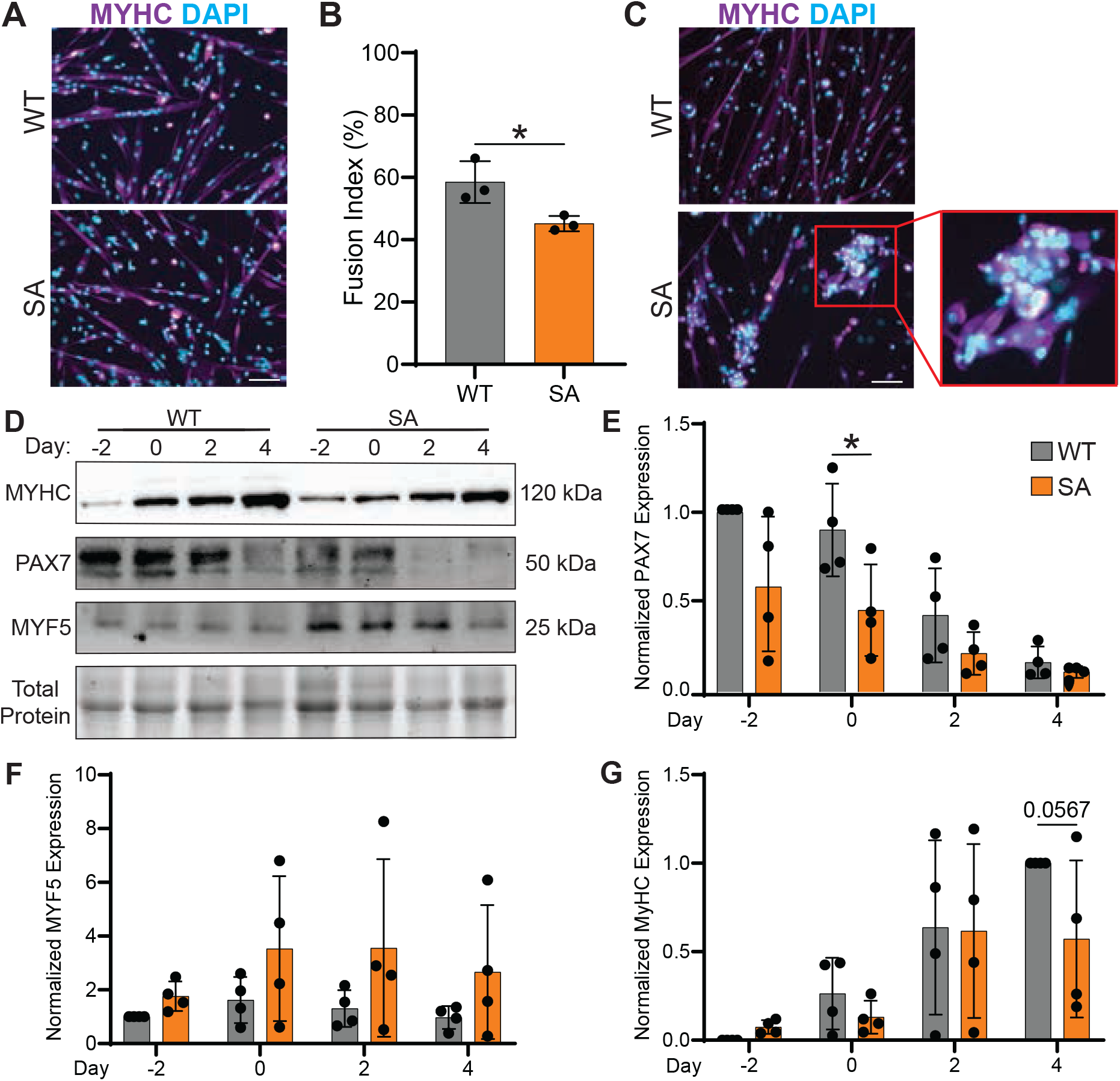
Inhibition of CARM1 phosphorylation at Ser572 impairs myogenic differentiation *in vitro*. CARM1 WT and SA primary myoblasts were differentiated for four days and assessed by immunostaining and immunoblotting. Cells were fixed at **A)** day 2 and **C)** day 4 and immunostained with anti-MYHC antibody (magenta) and nuclei were counterstained with DAPI (blue). **B)** Fusion index was quantified at day 2 of differentiation. **D)** Protein expression of PAX7, MYF5, and MYHC during myoblast differentiation at days -2, 0, 2, and 4 as determined by immunoblot analysis. Quantification of **E)** PAX7 and **F)** MYF5 protein abundance, normalized to total protein and expressed relative to WT day -2. **G)** MYHC protein abundance, normalized to total protein and expressed relative to WT day 4. Data are presented as mean ± SD; n = 3 biological replicates per genotype; *p ≤ 0.05; **B)** Students t-test; **E**, **F,** and **G)** two-way ANOVA with Fisher’s LSD post hoc test. Scale bars represent 100 µm.

To further characterize the myogenic state of cells during differentiation, we assessed the expression of myogenic proteins PAX7, MYF5, and MYHC. Notably, at the onset of differentiation (day 0), expression of the satellite stemness marker PAX7 in CARM1 SA myoblasts was reduced to 48.8% of wild-type levels (Fig. 2, D and E; p = 0.0491). Moreover, analysis of PAX7 expression across the differentiation time course revealed a significant effect of genotype, with CARM1 SA cells consistently expressing lower levels of PAX7 than wild-type cells (Fig. 2, D and E; two-way ANOVA, p = 0.0345 for genotype). A trend toward increased MYF5 protein abundance was observed in CARM1 SA cells; however, no statistically significant differences were detected between genotypes (Fig. 2, D and F). Additionally, CARM1 SA cells exhibited a trend toward reduced expression of the terminal differentiation marker MYHC at four days post-differentiation, reaching 57.18% of wild-type levels (Fig. 2, D and G; p = 0.0567). Together, these findings demonstrate that inhibition of CARM1 phosphorylation at Ser572 disrupts normal myogenic progression *in vitro*. Constitutive phospho-inhibition of CARM1 is associated with reduced expression of the satellite cell marker PAX7, impaired myoblast fusion and differentiation, and decreased expression of MYHC, supporting an important role for Ser572 phosphorylation during myogenic differentiation.

### Transcriptomic analyses reveal enhanced differentiation kinetics in CARM1 SA myoblasts

PAX7 is a master regulator of satellite cell identity and controls the expression of gene programs required for myogenic progression and differentiation (von Maltzahn et al., 2013). Given the reduced expression of PAX7 in CARM1 SA myoblasts, we next investigated whether loss of CARM1 Ser572 phosphorylation alters transcriptional programs during myogenic differentiation using RNA-sequencing. Under proliferative growth conditions, only 13 genes were differentially expressed between CARM1 SA and wild-type myoblasts (adjusted p < 0.05, absolute log_2_ fold change > 1), indicating that inhibition of CARM1 phosphorylation has minimal effects on the transcriptome of undifferentiated cells (Fig. S2 A; Table S3).

In contrast, 531 genes were differentially expressed at the onset of myogenic differentiation, demonstrating that transcriptional differences between genotypes become pronounced following differentiation induction (Fig. S2 B; Table S3). Functional enrichment analysis of these differentially expressed genes identified an upregulation of gene ontology (GO) terms associated with muscle organ development (GO: 0007517) and tissue development (GO: 0009888) in CARM1 SA cells (Fig. 3, A and B; Table S4). Consistent with these findings, inferred transcription factor activity analysis revealed increased activity of MYF5 and MYOD1, which promote myogenic commitment, as well as MYOG, a key regulator of terminal differentiation (Fig. 3 C; Table S5) (Zammit, 2017). Together, these data suggest that CARM1 SA myoblasts exhibit an enhanced propensity to activate myogenic differentiation programs.

**Figure 3.**
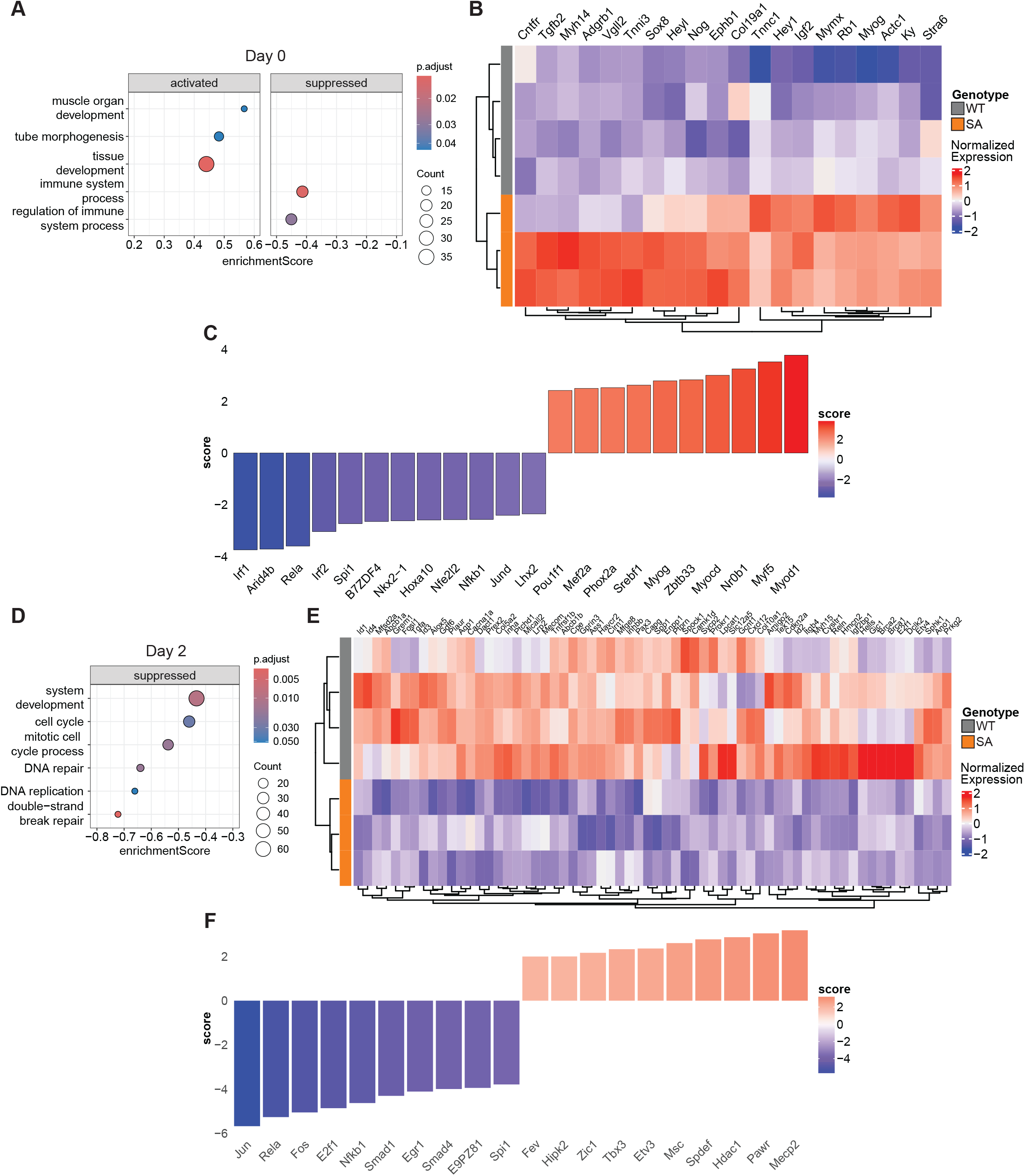
RNA sequencing reveals enrichment of myogenic differentiation pathways in CARM1 SA myoblasts. Bulk RNA-sequencing was performed on CARM1 WT and SA myoblasts at (**A-C**) day 0 and (**D-F**) day 2 of differentiation. **A**, **D**) Dot plots representing significantly enriched gene ontology (GO) terms identified from differentially expressed genes (DEGs; adjusted p-value < 0.05) in CARM1 SA relative to WT cells. **B, E)** Heat map of DEGs associated with the GO terms **B)** muscle organ development and **E)** system development. **C, F)** Inferred transcription factor activity analysis at **C)** day 0 and **F)** day 2 of differentiation. Transcription factor enrichment scores were calculated using a univariate linear model. Data are derived from n = 4 WT and 3 SA biological replicates.

To determine whether these transcriptional differences extended into differentiation, we compared the transcriptomes of CARM1 SA and wild-type cells two days after differentiation induction (Fig. S2 C; Table S3). At this time point, genes associated with cell cycle regulation (GO: 0007049) and system development (GO: 0048731) were significantly downregulated in CARM1 SA cells (Fig. 3, D and E; Table S4). Closer examination of genes within the system development category revealed decreased expression of factors known to restrain myogenic differentiation. Notably, expression of the *Id* family of transcriptional regulators, which function as inhibitors of myoblast differentiation, was reduced in CARM1 SA cells (Fig. 3 E) (Lingbeck et al., 2008). Since permanent cell cycle exit is a hallmark of terminal differentiation, the coordinated downregulation of cell cycle regulatory genes suggests that CARM1 SA cells transition more rapidly toward a differentiated, post-mitotic state (Holtzer et al., 1957). The advanced differentiation of CARM1 SA myoblasts is further supported by inferred transcription factor activity analysis, which identified reduced Fos transcription factor activity in CARM1 SA cells (Fig. 3 F; Table S5). Members of the Fos transcription factor family are associated with satellite cell activation, proliferation, and early stages of myogenic progression, and their activity typically declines as differentiation proceeds (Almada et al., 2021; Barutcu et al., 2022). Accordingly, reduced Fos transcriptional activity in CARM1 SA cells is consistent with progression beyond the early proliferative stages of myogenesis. Collectively, these transcriptomic analyses indicate that loss of CARM1 Ser572 phosphorylation accelerates activation of the myogenic differentiation program and promotes premature cell cycle withdrawal. However, despite acquiring a transcriptional profile characteristic of advanced differentiation, CARM1 SA cells exhibited impaired myoblast fusion and abnormal myotube formation (Fig. 2 A), suggesting that premature progression through the differentiation program ultimately compromises efficient myogenesis.

### Abrogation of CARM1 Ser572 phosphorylation induces precocious commitment and impairs self-renewal in satellite cells

Given our *in vitro* findings that inhibition of CARM1 Ser572 phosphorylation alters myogenic differentiation, we next sought to determine whether this phenotype originates at the level of satellite cell fate decisions. To directly monitor satellite cell activation, commitment, differentiation, and self-renewal, we employed an *ex vivo* single myofiber culture assay and assessed satellite cell identity by immunofluorescence staining for PAX7, MYOD, and MYOG.

During early activation (24 h), 66.9% of CARM1 SA satellite cells exhibited a committed myoblast identity (PAX7+/MYOD+) compared with 37.3% of wild-type cells, representing a 1.8-fold increase in the proportion of committed myogenic progenitors (Fig. 4, A and B; p = 0.0122). By 48 h, however, no differences in the PAX7+/MYOD+ population were observed between genotypes (Fig. 4, D and E). These data indicate that loss of CARM1 Ser572 phosphorylation accelerates the timing of satellite cell commitment without affecting the overall extent of commitment (Fig. 4, A-F).

**Figure 4.**
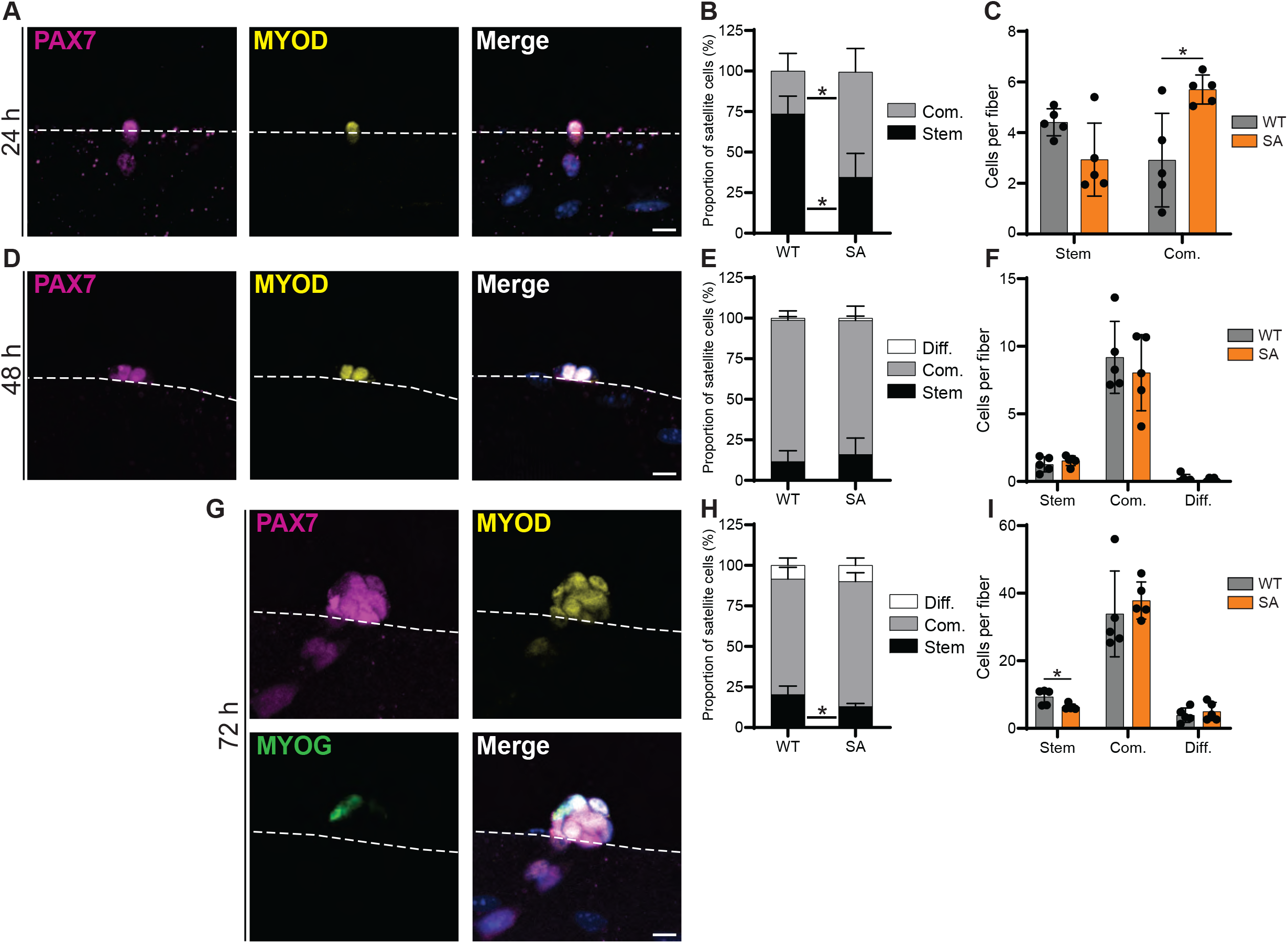
Inhibition of CARM1 phosphorylation induces precocious commitment and impaired satellite cell self-renewal. Single myofibers isolated from EDL muscles of male CARM1 WT and SA mice were cultured for **A)** 24 h **D)** 48 h, and **G)** 72 h. Myofibers were immunostained for PAX7 (magenta), MYOD (yellow), and MYOG (green); and nuclei were counterstained with Hoechst (blue). Representative images are from CARM1 WT myofibers. Satellite cells were classified as stem (PAX7+ only), committed (Com.; PAX7+/MYOD+), or differentiating (Diff.; PAX7-). The percentage of each subpopulation was quantified at **B)** 24 h, **E)** 48 h, and H) 72 h. The number of cells within each subpopulation was normalized to the number of myofibers analyzed at **C)** 24 h, **F)** 48 h, and **I)** 72 h. Data are presented as mean ± SD; n = 5 biological replicates per genotype; Student’s t-test; *p ≤ 0.05. Scale bars represent 10 µm.

At 72 h post-isolation, three populations of myogenic progenitors were identified: self-renewing satellite cells (PAX7+ only), committed myoblasts (PAX7+/MYOD+), and differentiating myocytes (PAX7-) (Zammit et al., 2004). No differences were observed in the proportions of committed myoblasts or differentiating myocytes between CARM1 SA and wild-type cultures (Fig. 4, G and H). In contrast, the self-renewing satellite cell population was reduced by 36.01% in CARM1 SA myofiber cultures compared with wild-type controls (Fig. 4, G and H; p = 0.0176). Together, these findings reveal two major consequences of CARM1 Ser572 phospho-inhibition. First, satellite cells enter the myogenic program more rapidly during the early stages of activation, consistent with the accelerated differentiation phenotype observed *in vitro*. Second, this altered fate trajectory reduces the pool of self-renewing satellite cells, suggesting that CARM1 phosphorylation at Ser572 is required to maintain an appropriate balance between commitment and self-renewal during satellite cell activation.

### Abrogation of CARM1 Ser572 phosphorylation impedes satellite cell maintenance following injury-induced regeneration

The reduction in self-renewing satellite cells observed in the single myofiber assay prompted us to examine whether inhibition of CARM1 Ser572 phosphorylation affects maintenance of the satellite cell pool *in vivo*. We first examined muscle morphology and satellite cell content from three-month-old adult mice. Histological analysis of tibialis anterior (TA) muscles revealed no differences in muscle morphology between wild-type and CARM1 SA mice, nor were differences observed in muscle mass (Fig. 5, A and B) or muscle fiber size (Fig. S3, A-C). Similarly, quantification of PAX7+ satellite cells in TA cross sections revealed no differences in satellite cell content (Fig. 5, G and H).

**Figure 5.**
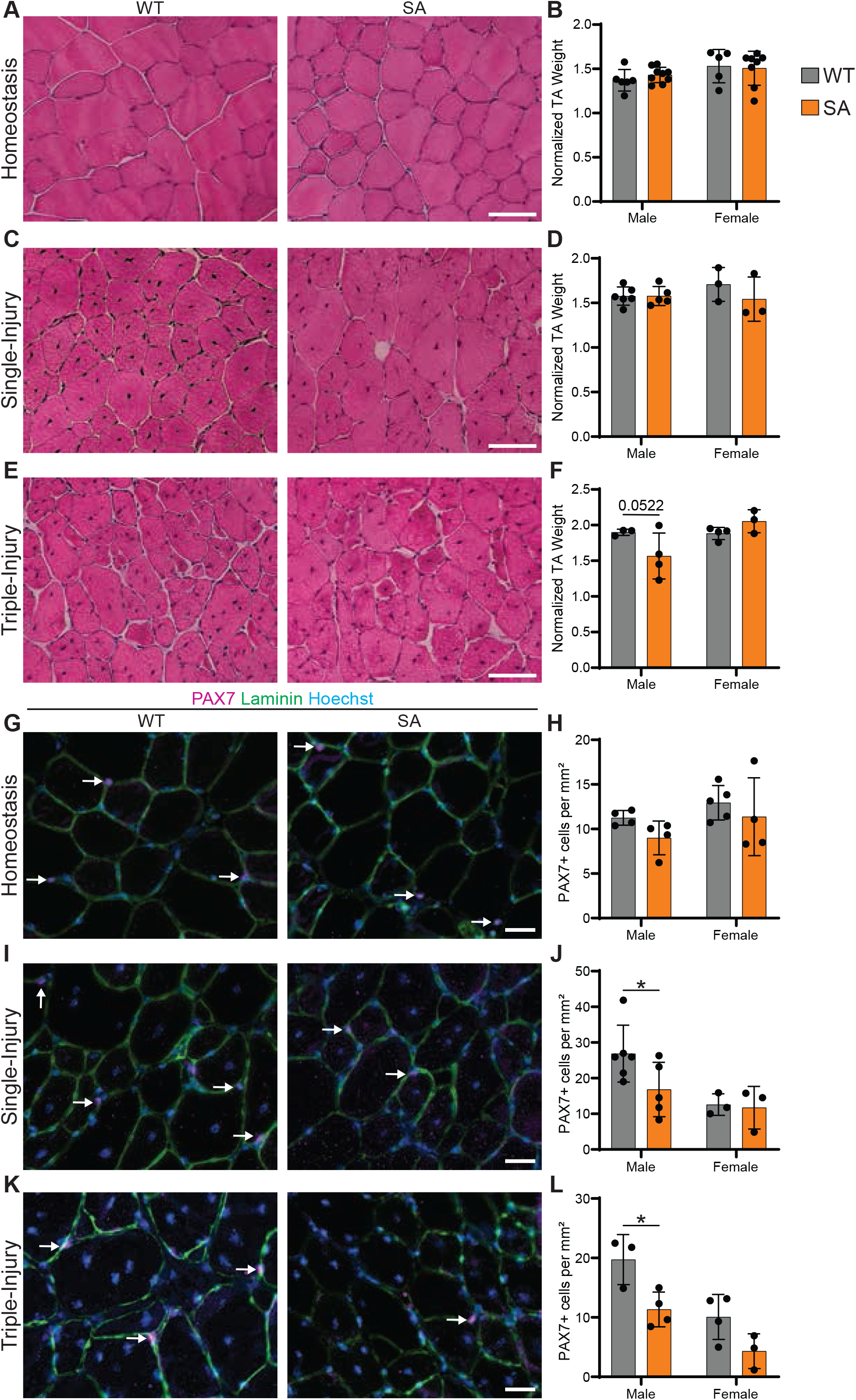
Inhibition of CARM1 phosphorylation at Ser572 impairs satellite cell self-renewal following *in vivo* muscle injury. Tibialis anterior (TA) muscles from CARM1 WT and SA mice were assessed at homeostasis and following regeneration induced by either one or three rounds of intramuscular injury. Representative H&E stained TA cross-sections from **A)** homeostatic, **C)** single-injury, and **E)** triple-injury muscles. Muscle mass normalized to total body weight for **B**) homeostatic, **D**) single-injury and **F**) triple-injury muscles. TA cross-sections from **G)** homeostatic, **I)** single-injury, and **K)** triple-injury mice were immunostained for PAX7 (magenta), laminin (green), and nuclei were counterstained with Hoechst (blue). Quantification of PAX7+ satellite cells normalized to tissue area is shown for **H)** homeostatic, **J)** single-injury, and **L)** triple-injury muscles. Data are presented as mean ± SD; n = 3-9 biological replicates per sex and genotype as indicated; two-way ANOVA with Fisher’s LSD post hoc test; *p ≤ 0.05. Scale bars represent 50 µm (**A, C, E**) and 20 µm (**G, I, K**).

We next examined whether the impaired differentiation phenotype observed *in vitro* was reflected in regenerative capacity following acute muscle injury. Despite the defects observed *in vitro*, injured CARM1 SA TA muscles displayed normal muscle morphology, muscle mass, and muscle fiber size compared with wild-type controls after 21 days of regeneration (Fig. 5, C and D; and Fig. S3, D-F). However, analysis of satellite cell content revealed a significant reduction in regenerated muscles from male CARM1 SA mice, reaching 58.2% of wild-type levels (Fig. 5, I and J; p = 0.0342). However, no differences were observed between female CARM1 WT and SA mice (Fig. 5, I and J). These findings suggest that inhibition of CARM1 Ser572 phosphorylation does not impair muscle regeneration *in vivo* but compromises maintenance of the satellite cell pool following regeneration, and this difference may be sex-dependent.

To address long-term satellite cell maintenance, we subjected mice to repeated rounds of muscle injury. Following three rounds of regeneration, TA muscle mass in male CARM1 SA mice was reduced to 82.5% of wild-type levels, although this difference did not reach statistical significance (Fig. 5 F; p = 0.0522). No differences in muscle mass were observed between female mice (Fig. 5 F). Notably, two-way ANOVA identified a significant interaction between sex and genotype (p = 0.0398), indicating sex-dependent effects of CARM1 Ser572 phospho-inhibition on long-term muscle recovery. Muscle fiber size showed evidence of a similar sex-dependent pattern, although the interaction effect did not reach statistical significance (Fig. S3, G-I; p = 0.1345). Analysis of satellite cell content revealed a pronounced depletion of the satellite cell pool following repeated injury. Two-way ANOVA identified significant effects of both genotype and sex (Fig. 5, K and L; p = 0.0037 and p = 0.0017, respectively). When analyzed by sex, satellite cell content was reduced to 57.51% of wild-type levels in males (p = 0.0103) and 42.75% of wild-type levels in females (Fig. 5 L; p = 0.0553). Collectively, these *in vivo* studies demonstrate that CARM1 Ser572 phosphorylation is dispensable for muscle development and acute regeneration but is required for long-term maintenance of the satellite cell pool. The progressive depletion of satellite cells following regenerative challenge supports a role for CARM1 phosphorylation in promoting satellite cell self-renewal and sustaining regenerative capacity over time.

### Transcriptional analysis reveals impaired self-renewal of satellite cells during regenerative myogenesis

To identify the altered biological processes underlying the loss of satellite cell self-renewal, we performed RNA sequencing on satellite cells isolated under homeostatic and regenerative (three days post-injury; 3 DPI) conditions (Fig. S4, A and B). The 3 DPI time point was selected because it contains a distinct self-renewing satellite cell subpopulation (Okafor et al., 2023). Under homeostatic conditions, only 43 genes were differentially expressed between wild-type and CARM1 SA satellite cells (Fig. 6 A; Table S3; adjusted p-value < 0.05), and no significantly enriched biological processes were identified by GO analysis. In contrast, 331 genes were differentially expressed at 3 DPI (Fig. 6 B; Table S3; adjusted p-value < 0.05), indicating that the transcriptional consequences of CARM1 Ser572 phospho-inhibition become most apparent during regeneration.

**Figure 6.**
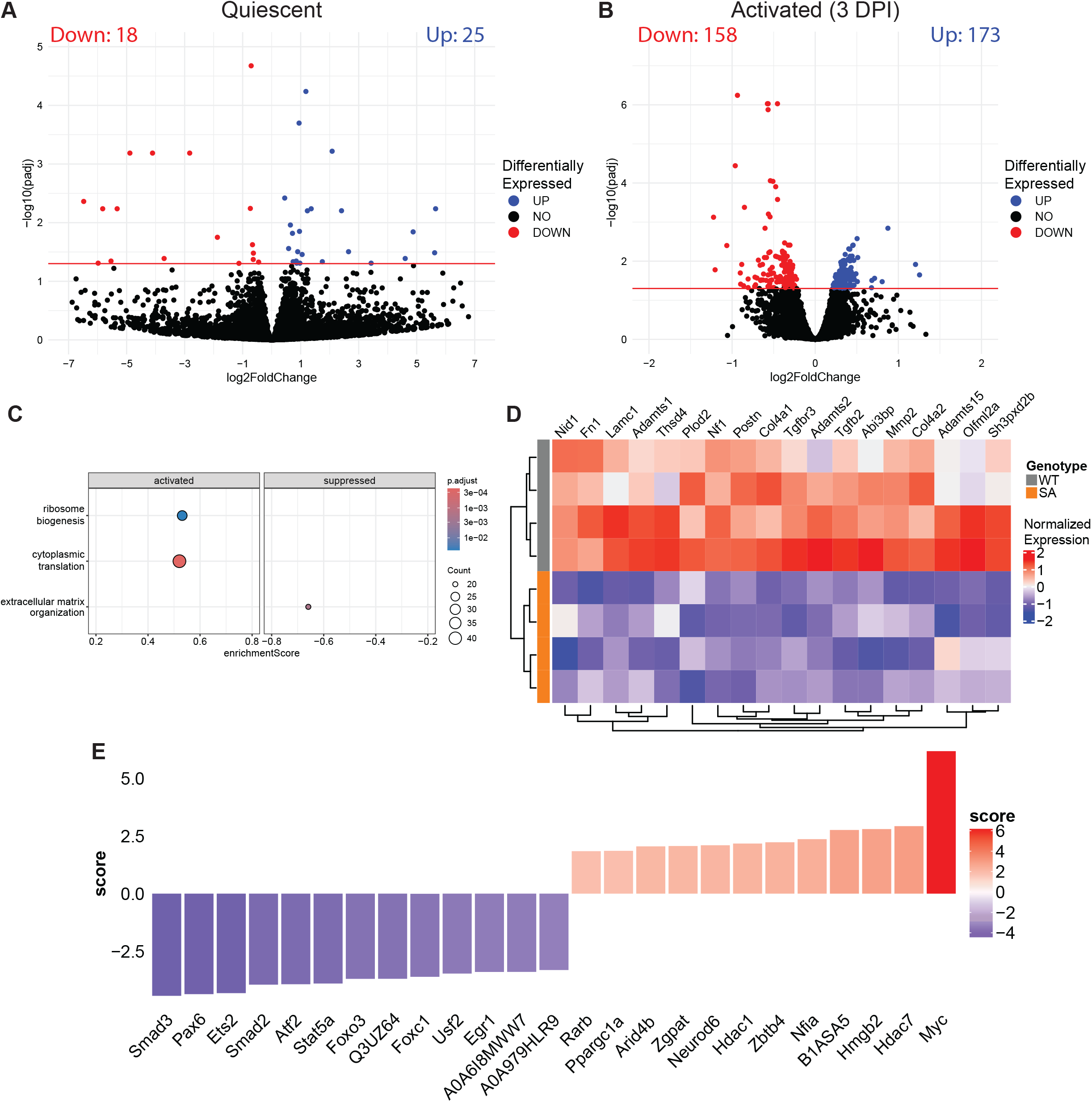
RNA sequencing of activated satellite cells identifies transcriptional alterations associated with impaired satellite cell self-renewal. Bulk RNA sequencing was performed on freshly isolated satellite cells from **A)** homeostatic (representing quiescent satellite cells) and **B)** regenerating hindlimb muscle at 3 days post-injury (3 DPI; representing activated satellite cells). Volcano plots show differentially expressed genes (DEGs; adjusted p-value < 0.05) in CARM1 SA relative to WT satellite cells. **C)** Dotplot representing enriched gene ontology (GO) terms identified from DEGs in activated satellite cells. **D)** Heatmap of DEGs associated with the extracellular matrix organization GO term. **E)** Inferred transcription factor activity analysis based on DEGs identified in activated satellite cells. For **A**, n = 5 WT (3 male, 2 female) and 4 CARM1 SA (2 male, 2 female). For **B-E**, n = 4 for each genotype (2 male, 2 female).

Gene set enrichment analysis identified extracellular matrix (ECM) organization (GO:0030198) as one of the most significantly downregulated pathways in activated CARM1 SA satellite cells (Fig. 6 C; Table S4). During muscle regeneration, reciprocal interactions between satellite cells and ECM are essential for satellite cell expansion and self-renewal, we therefore examined the associated genes to gain better insight into what pathways are affected (Baghdadi et al., 2018; Bentzinger et al., 2013; Chrysostomou and Mourikis, 2024; Goetsch et al., 2003; Rayagiri et al., 2018). Among the downregulated ECM-associated genes was *Fn1*, which encodes fibronectin (Fig. 6 D). Fibronectin deposition by satellite cells promotes expansion of Pax7-positive progenitors and is required for effective satellite cell self-renewal during muscle regeneration (Bentzinger et al., 2013). We also observed reduced expression of *Tgfbr3*, which encodes betaglycan (Fig. 6 D), a recently identified marker of the self-renewing satellite cell subpopulation (Okafor et al., 2023). Together, these transcriptional changes provide direct evidence that loss of CARM1 Ser572 phosphorylation disrupts molecular pathways associated with satellite cell self-renewal.

Because TGFβ signaling promotes satellite cell self-renewal through SMAD-mediated transcriptional programs (Okafor et al., 2023), we next examined inferred transcription factor activity. Consistent with reduced *Tgfbr3* expression, inferred transcription factor analysis identified diminished SMAD2 and SMAD3 activity in activated CARM1 SA satellite cells (Fig. 6 E; Table S5) (Okafor et al., 2023). SMAD4 activity was also reduced, although it was not among the most significantly altered transcription factors (Table S5). Notably, reduced SMAD4 activity was also observed during *in vitro* myoblast differentiation (Fig. 3 F), suggesting that impaired TGFβ-SMAD signaling represents a common consequence of CARM1 Ser572 phospho-inhibition. The transcription factor with the most significantly elevated activity was MYC (Fig. 6 E). MYC is induced during early satellite cell activation and promotes chromatin remodeling required for entry into the myogenic program (He et al., 2021). Elevated MYC activity is therefore consistent with enhanced satellite cell activation and may contribute to the accelerated commitment phenotype observed in CARM1 SA satellite cells.

Collectively, these transcriptomic analyses indicate that CARM1 Ser572 phospho-inhibition disrupts transcriptional programs that support satellite cell self-renewal during regeneration. Downregulation of ECM-associated genes, including *Fn1* and *Tgfbr3*, together with impaired TGFβ-SMAD signaling, are consistent with depletion of the self-renewing satellite cell population. Concurrently, increased MYC activity supports an accelerated activation phenotype, suggesting that enhanced commitment occurs at the expense of self-renewal, ultimately contributing to the progressive loss of satellite cells observed following regeneration.

## Discussion

Skeletal muscle maintenance and regeneration rely on the precise regulation of satellite cell fate decisions (Relaix et al., 2021). To determine the physiological role of p38γ-mediated phosphorylation of CARM1 at Ser572, we generated mice carrying a non-phosphorylatable serine-to-alanine mutation. Myogenic progenitors harboring the CARM1 SA mutation underwent precocious myogenic commitment, resulting in a reduction in the self-renewing satellite cell population. Notably, in single myofiber culture assays, we did not observe differences in the total number of committed satellite cells or those undergoing terminal differentiation at later time points. These findings indicate that accelerated commitment primarily reduces entry into the self-renewal pathway rather than altering the extent of terminal differentiation.

Despite the reduction in satellite cell content following regeneration *in vivo*, muscle repair was largely preserved after a single injury. In contrast, deletion of p38γ impairs both satellite cell maintenance and regenerative capacity (Chang et al., 2018). Together, these findings suggest that loss of CARM1 Ser572 phosphorylation accounts for the self-renewal phenotype of p38γ deficiency but is insufficient to explain the regeneration defects observed following p38γ deletion, which likely arise from additional downstream targets of p38γ signaling (Gillespie et al., 2009).

Interestingly, the defects in myogenic differentiation observed *in vitro* did not translate into overt impairments in muscle regeneration following acute injury *in vivo*. Primary CARM1 SA myoblasts cultured *in vitro* exhibited reduced fusion efficiency, abnormal myotube morphology, and decreased expression of differentiation-associated markers, indicating impaired progression through the myogenic program. In contrast, muscle regeneration proceeded normally *in vivo,* with no significant deficits in myogenesis detected following injury. This discrepancy likely reflects the distinct biological contexts of the *in vitro* and *in vivo* environments. Regeneration *in vivo* occurs within a highly coordinated niche composed of inflammatory cells, fibro-adipogenic progenitors, endothelial cells, extracellular matrix components, and mechanical cues that collectively promote efficient myogenesis and can compensate for intrinsic cellular defects. In contrast, cultured myoblasts differentiate in the absence of many of these regulatory inputs, potentially altering the presentation of phenotypes that are present during regeneration. Thus, both the *in vitro* differentiation defects and the *in vivo* self-renewal phenotype may arise from a common underlying mechanism: premature entry into the myogenic program at the expense of stem cell maintenance.

Transcriptomic analysis identified impaired extracellular matrix remodeling and TGFβ signaling as potential mechanisms underlying the self-renewal defect. Reduced expression of *Fn1* and *Tgfbr3*, together with diminished SMAD2/3 activity, suggests that loss of CARM1 Ser572 phosphorylation disrupts pathways that normally support satellite cell maintenance and expansion. However, the molecular link between CARM1 Ser572 phosphorylation and these self-renewal programs remains unclear. The observed reduction in *Tgfbr3* expression and SMAD activity raises the possibility that altered commitment disrupts the acquisition or maintenance of the recently described TGFBR3+ self-renewing satellite cell state. Future studies examining CARM1-dependent chromatin landscapes may determine whether components of the TGFβ-SMAD pathway are direct targets of CARM1-mediated epigenetic regulation or whether their dysregulation arises secondarily as a consequence of altered cell fate progression (Okafor et al., 2023; Zheng et al., 2021).

The context-dependent nature of the CARM1 SA phenotype in satellite cells *in vivo* suggests that CARM1 Ser572 phosphorylation acts primarily during periods of cell-fate transition. Satellite cells in uninjured muscle remain largely quiescent and exhibit minimal transcriptional perturbation in the absence of Ser572 phosphorylation. In contrast, substantial phenotypic and transcriptional differences emerged following activation, when satellite cells must repeatedly choose between commitment and self-renewal.

Our findings are consistent with a model in which phosphorylation regulates the timing rather than efficiency of CARM1 activity. CARM1 is required for myogenic commitment; however, phosphorylation at Ser572 restricts its ability to promote commitment by limiting its nuclear translocation. Consequently, preventing phosphorylation does not inhibit CARM1 function but instead removes a regulatory mechanism that under normal conditions would restrain commitment. Loss of Ser572 phosphorylation therefore accelerates progression through the commitment phase, leading to premature depletion of self-renewing progenitors.

We also demonstrated that CARM1 expression, activity, and localization undergo dynamic changes during myogenic differentiation. CARM1 expression increased during satellite cell activation, whereas protein abundance, methyltransferase activity, and nuclear localization declined as differentiation progressed. A potential mechanism underlying the decline in CARM1 abundance during differentiation is regulation by the miR-181 family, which suppresses CARM1 expression during stem cell differentiation and is induced during myogenesis (Naguibneva et al., 2006; Xu et al., 2013).

One limitation of the present study is that the Ser-to-Ala mutation was introduced into both CARM1 isoforms. Although previous studies identified phosphorylation-dependent regulation of the CARM1Δ15-PAX7 axis, the potential contribution of full-length CARM1 cannot be excluded (Chang et al., 2018). Some aspects of the observed phenotype may arise from phosphorylation-dependent regulation of additional targets. Development of isoform-specific phospho-mutant models will be required to fully resolve the relative contributions of individual CARM1 isoforms in satellite cell fate determination.

Of note, we observed sex-dependent effects of the CARM1 SA mutation. Following a single injury, reduced satellite cell content was detected in male but not female mice, whereas repeated injury depleted the satellite cell pool in both sexes. Although the mechanisms underlying these differences remain unclear, CARM1 functions as a coactivator of both androgen and estrogen receptors, and satellite cells exhibit distinct responses to androgen and estrogen signaling (Chen et al., 1999; Collins et al., 2019; Seko et al., 2020; Sinha-Hikim et al., 2004). Consequently, differential hormone receptor signaling may contribute to the sex-specific effects of the CARM1 SA mutation.

Our findings also raise the possibility that modulation of CARM1 Ser572 phosphorylation could be beneficial in disease contexts characterized by impaired satellite cell commitment, such as in DMD. Satellite cells in DMD exhibit a shift toward self-renewal at the expense of commitment, as evidenced by satellite cell hyperplasia and recent studies demonstrating a stalled commitment program that impairs regenerative myogenesis (Esper et al., 2025; Granet et al., 2025). Previous work showed that disruption of dystrophin-associated signaling leads to dysregulated p38γ activity and hyperphosphorylation of CARM1 at Ser572, resulting in impaired satellite cell commitment (Chang et al., 2018). Together with our current study, these findings suggest that optimal muscle regeneration requires a finely tuned level of CARM1 phosphorylation rather than complete activation or inhibition of the pathway. While preventing CARM1 phosphorylation may be detrimental in healthy muscle by inducing stem cell depletion, it will be important to determine if reducing excessive Ser572 CARM1 phosphorylation within a DMD context could restore commitment and improve regenerative capacity. These observations highlight the importance of temporal and quantitative control of CARM1 activity as a determinant of satellite cell fate.

Collectively, our findings identify CARM1 Ser572 phosphorylation as a key regulator of satellite cell fate decisions during regenerative myogenesis. By restraining the rate of satellite cell commitment, phosphorylation at Ser572 promotes entry into the self-renewal pathway and supports long-term maintenance of the satellite cell pool. These results highlight the importance of balancing commitment and self-renewal during muscle regeneration and establish CARM1 Ser572 phosphorylation as a molecular mechanism that maintains this balance.

## Materials and Methods

### Generation of CARM1 Ser572/595 phosphorylation-inhibitory mutant mice

Ser572 (CARM1Δ15) / Ser595 (full-length CARM1) phosphorylation-inhibitory mutant mice (CARM1 SA) were generated by the Model Production Core at The Centre for Phenogenomics using CRISPR-Cas9-mediated genome editing in C57BL/6 mice. Murine zygotes were electroporated with Cas9, a single guide RNA (sgRNA; GGTCGGGAT), and a single-stranded DNA repair template encoding a Ser-to-Ala substitution (Fig. S1). The repair template induced three nucleotide changes to *Carm1*, affecting both isoforms: two substitutions within the TCG codon encoding Ser572/595 (c.1897/1828T>G and c.1899/1830G>C) generated an Ala572/595 residue, while a third silent substitution (c.1896/1827C>T) was incorporated within the PAM sequence to prevent re-cutting by Cas9. Genome editing was confirmed by PCR amplification of the targeted locus followed by Sanger sequencing.

Founder CARM1 SA mice were crossed onto the C57BL/10 background and backcrossed for six generations. Heterozygous offspring were subsequently intercrossed to establish the colony. Wild-type and homozygous CARM1 SA mice were used for all experiments. Genotypes were originally confirmed by endpoint PCR and subsequently by probe-based qPCR (Table S1). All animal procedures were approved by the McGill University Animal Care Committee (MCGL-8124). Animal housing, husbandry, and experimental procedures were conducted in accordance with the guidelines of the McGill University Animal Compliance Office and the Canadian Council on Animal Care.

### Primary myoblast isolation, culture, and differentiation

Primary myoblasts were generated from CARM1 SA mice by magnetic cell separation (MACS), as previously described (Filippelli et al., 2021). Briefly, hindlimb muscles were mechanically dissociated in a collagenase B (Roche) and dispase II (Roche) solution using a gentleMACS Octo Dissociator (Miltenyi Biotec). Cell suspensions underwent negative selection using the Satellite Cell Isolation Kit (Miltenyi Biotec) followed by positive selection with anti-integrin α7 MicroBeads (Miltenyi Biotec). Purified cells were seeded onto plates coated with type I collagen. Primary myoblasts were maintained in growth media (GM) consisting of Ham’s F-10 supplemented with 20% FBS, 1X penicillin/streptomycin, and 25 ng/mL of basic fibroblast growth factor (Thermo Fisher) at 37°C in a humidified incubator containing 5% CO_2_. Differentiation was induced at confluence by replacing GM with differentiation medium (DM) containing 45% F-10, 45% DMEM, and 10% horse serum, as previously described (VanGenderen et al., 2022).

### Immunocytochemistry

Immunofluorescence (IF) staining of cells was performed as previously described (VanGenderen et al., 2022). Cells were fixed in 4% paraformaldehyde (PFA), permeabilized, blocked, and incubated with primary antibodies overnight at 4°C (Table S2). Fluorescent secondary antibodies were applied for 1 h, nuclei were counterstained with Hoechst, and samples were mounted using ProLong Gold Antifade reagent (Thermo Fisher). Samples were imaged on a Zeiss Axio Observer 7. For *in vitro* differentiation experiments, fusion index was calculated with the formula: 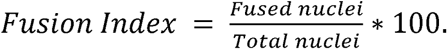

### Western blotting

Western blotting was performed as previously described (VanGenderen et al., 2022). Cells were lysed in lysis buffer (50 mM Tris HCl pH 7.5, 150 mM NaCl, 0.5% Triton X-100, 0.5 mM EDTA, 1X protease (Nacalai), and 1X phosphatase inhibitors (Nacalai)). Proteins were resolved by SDS-PAGE on gels containing 0.5% 2,2,2-trichloroethanol (TCE), and total protein loading was visualized by UV activation using a ChemiDoc Imaging System (Bio-Rad). Proteins were subsequently transferred to polyvinylidene fluoride (PVDF) membranes using a Trans-Blot Turbo Transfer System (Bio-Rad). Membranes were blocked, incubated overnight with primary antibodies (Table S2), and probed with HRP-conjugated secondary antibodies (Bio-Rad; 1:10,000). Protein detection was performed using enhanced chemiluminescence (Bio-Rad) and visualized on a ChemiDoc Imaging System. Protein abundance was quantified using Image Lab v6.1 software (Bio-Rad) and normalized to total protein loading.

### Single myofiber isolation and staining

Single myofiber isolation and staining were performed as described previously (Granet et al., 2025). Briefly, extensor digitorum longus (EDL) muscles were dissected from tendon to tendon and digested with collagenase type 1. Individual myofibers were isolated and maintained in myofiber culture medium (DMEM supplemented with 20% FBS, 1% chick embryo extract, 1% P/S, and 125 ng/mL bFGF). Myofibers were fixed in PFA, permeabilized, blocked, and stained overnight with primary antibodies (Table S2), followed by fluorescent secondary antibodies and Hoechst nuclear stain. Myofibers were mounted with ProLong Gold Antifade mounting media. Fibers were imaged and quantified using an EVOS M5000 microscope (Thermo Fisher).

### Satellite cell isolation

Homeostatic and activated satellite cells were isolated from 9-to 13-week-old mice by fluorescence activated cell sorting (FACS) as previously described (Pasut et al., 2012). Hindlimb muscles were dissected and enzymatically dissociated in a collagenase B and dispase II solution using a gentleMACS OctoDissociator. Cell suspensions were filtered, subjected to red blood cell lysis (Sigma), and stained with antibodies (Table S2) (Pasut et al., 2012). Satellite cells were isolated using a FACSAria III cell sorter (BD Biosciences) and identified as ITGA7+/VCAM1+/Lin-(CD31, CD45, CD11b, and Sca1) cells. For activated satellite cell isolation, muscle injury was induced by intramuscular injection of cardiotoxin (CTX, Latoxan) into the tibialis anterior (TA, 30 μL, 10 μM) and gastrocnemius (50 μL, 10 μM). Satellite cells were isolated 3 days post-injury (DPI).

### RNA sequencing and bioinformatic analysis

#### Primary myoblasts

Total RNA was isolated from cultured myoblasts using Trizol and purified with the Qiagen RNeasy Kit as previously described (VanGenderen et al., 2022). Library preparation was performed using the TruSeq Library Prep Kit (Illumina) and sequenced on a NextSeq 500 (Illumina) at the Institute for Research in Immunology and Cancer (IRIC), Université de Montréal.

#### Homeostatic satellite cells

RNA isolation, cDNA synthesis, and amplification were performed using the SMART-Seq HT Kit (Takara) according to the manufacturer’s instructions. Briefly, 2,000 FACS-isolated satellite cells were directly lysed and subjected to cDNA synthesis and amplification. Following purification with AMPure XP beads (Beckman Coulter), cDNA concentration was assessed using a Quant-iT PicoGreen dsDNA Assay Kit (Life Technologies). Sequencing libraries were prepared using the Nextera XT Library Preparation Kit (Illumina) and sequenced on a NextSeq 500 platform.

#### Activated satellite cells

Total RNA was isolated from activated satellite cells using the Direct-zol RNA Kit (Zymo Research) according to the manufacturer’s instructions and quantified using a Qubit RNA HS Assay Kit (Thermo Fisher). Following poly (A) enrichment (New England Biolabs), RNA-seq libraries were generated using the NEBNext Ultra II RNA Library Prep Kit for Illumina (New England Biolabs) according to the manufacturer’s protocol. Sequencing was performed on a NextSeq 2000 platform (Illumina) at IRIC.

#### Bioinformatics analysis

Sequencing reads were aligned to the murine GRCm38 reference genome using STAR within the GenPipes v4.0 pipeline (Bourgey et al., 2019; Dobin et al., 2013; Putri et al., 2022). Raw transcript counts were generated using HTSeq. Differential gene expression analysis was performed by DESeq2 (Love et al., 2014). For activated satellite cells, batch effects were corrected prior to DESeq2 analysis using the ComBat-seq function of the sva package (Leek et al., 2012). Gene set enrichment analysis was performed using the gseGO function of clusterProfiler on differentially expressed genes (DEGs) with an adjusted p-value of < 0.05 (Wu et al., 2021). Transcription factor activity was inferred using the decoupleR package (Badia-i-Mompel et al., 2022).

### Muscle injury, regeneration, and histology

Muscle regeneration was induced in 10 to 14-week-old mice by injection of 50 μL of 1.2% BaCl in saline into the tibialis anterior (TA) muscle. For single injury experiments, muscles were collected at 21 days post-injury (21 DPI). For repeated injury experiments, BaCl administration was performed three times, with each injury followed by a 21-day regeneration period. Muscles were harvested and snap-frozen in liquid nitrogen-cooled isopentane.

All histological analyses were performed on PFA-fixed 10 μm transverse cryosections of TA muscles. Hematoxylin and eosin staining was performed using standard procedures. For immunofluorescence, sections were fixed, subjected to antigen retrieval by citrate decrosslinking (Abcam), permeabilized, and blocked prior to overnight incubation with primary antibodies (Table S2). Fluorescent secondary antibodies and Hoechst nuclear stain were applied before mounting in ProLong Gold Antifade reagent (Thermo Fisher). Sections were imaged using a Zeiss Axio Observer 7 microscope, and PAX7+ cells were quantified using Fiji/ImageJ (version 1.54p) (Schindelin et al., 2012).

### Statistical analysis

Statistical analyses were performed using GraphPad Prism v10.2.0. Data are presented as mean ± standard deviation (SD). Statistical significance was determined by either two-tailed Student’s t-tests or two-way ANOVA followed by Fisher’s least significant difference (LSD) or Tukey’s multiple-comparison tests, as indicated. P-values for t-tests and post-hoc multiple comparisons are indicated as *p ≤ 0.05, **p ≤ 0.01, ***p ≤ 0.001.

## Supplemental Material

## Supplementary tables

**Table S1.** Primer and probe sequences used in this study.

**Table S2.** Antibodies used and applications.

**Table S3.** Differentially expressed genes identified between CARM1 WT and SA myoblasts and satellite cells (related to Fig. S2, A-C; and Fig. 6, A-B).

**Table S4.** Gene ontology enrichment of differentially expressed genes in CARM1 WT and SA myoblasts and satellite cells (related to Fig. 2, A and D; and Fig. 6 C).

**Table S5.** Inferred transcription factor activity in CARM1 WT and SA myoblasts and satellite cells (related to Fig. 3, C and F; and Fig. 6 E).

## Supplementary figures

**Figure S1.** CRISPR/Cas9-mediated mutagenesis of the CARM1 Ser572/595 residue.

**Figure S2.** Transcriptomic profiling of CARM1 WT and SA myoblasts during myogenic differentiation.

**Figure S3.** Muscle fiber cross-sectional area and size distribution in CARM1 wild-type and SA mice.

**Figure S4.** Principal component analysis of transcriptomic profiles from quiescent and activated satellite cells in CARM1 wild-type and SA mice.

## Data availability

RNA-sequencing datasets generated and analyzed in this study and corresponding to the experiments shown in Fig. 3 and 6 have been deposited in the NCBI Gene Expression Omnibus (GEO) under accession number GSE348227 and will be released upon publication. Reviewer access credentials are provided to the editors and reviewers. All other data supporting the findings of this study are available from the corresponding author upon reasonable request.

## Acknowledgments

We thank Camille Stegen and Julien Leconte of the Flow Cytometry Core Facility, Frédérique Dansereau of the Comparative Medicine and Animal Resources Centre, and Nicolas Audet of the Imaging and Molecular Biology Platform (IMBP), McGill University, for technical support and expertise. We also thank Raphaëlle Lambert of the Genomics Platform at the Institute for Research in Immunology and Cancer (IRIC), Université de Montréal, for RNA sequencing support. We thank Mark Bedford (MD Anderson) for kindly providing the anti-methyl-NFIB antibody. Additionally, we are grateful to Chang lab members Romina Filippelli, Tim Lorenz, and Alessio Cusmano for their assistance in experiments. C.A.V. is the recipient of Graduate Excellence Award, Gerald Clavet Fellowship in Medicine, Shuk-Tak Liang Fellowship in Medicine and David G Guthrie Fellowship in Medicine from McGill University, in addition to Fonds de recherche du Québec-Santé (FRQS) Bourses de doctorant en recherche (365505). N.C.C. is the recipient of FRQS Junior 1 and Junior 2 Chercheuse-Boursière awards (298146, 309857, and 366069). This work was supported by grants from the Canadian Institutes of Health Research (CIHR; 175431 and 180499) to N.C.C.

## Author contributions

Cordell A. VanGenderen: conceptualization, data curation, formal analysis, investigation, visualization, and writing - original draft, review, and editing. Rebecca Robertson: formal analysis, investigation, and writing - review and editing. Jules A. Granet: investigation. Ishtiaque Hossain: methodology. Ronit Mohapatra: investigation. John Saber: methodology. Michael A. Rudnicki: resources and supervision. William A. Pastor: conceptualization and supervision. Natasha C. Chang: conceptualization, formal analysis, funding acquisition, investigation, project administration, supervision, and writing - original draft, review, and editing.

