## Supplemental Information for "Phosphorylation of CARM1 at Ser572 preserves muscle stem cell self-renewal by restricting premature myogenic commitment"

**Supplementary tables:**

**Table S1.** Primer and probe sequences used in this study.

**Table S2.** Antibodies used and applications.

**Table S3.** Differentially expressed genes identified between CARM1 WT and SA myoblasts and satellite cells (related to Figures S2A-C and 6A-B).

**Table S4.** Gene ontology enrichment of differentially expressed genes in CARM1 WT and SA myoblasts and satellite cells (related to Figures 2A, 2D, and 6C).

**Table S5.** Inferred transcription factor activity in CARM1 WT and SA myoblasts and satellite cells (related to Figures 3C, 3F, 6E).

**Supplementary figures:**

**Figure S1.** CRISPR/Cas9-mediated mutagenesis of the CARM1 Ser572/595 residue.

**Figure S2.** Transcriptomic profiling of CARM1 WT and SA myoblasts during myogenic differentiation.

**Figure S3.** Muscle fiber cross-sectional area and size distribution in CARM1 wild-type and SA mice.

**Figure S4.** Principal component analysis of transcriptomic profiles from quiescent and activated satellite cells in CARM1 wild-type and SA mice.

**Table S1.** Primer and probe sequences used in this study.

| Application | Primer Name | Sequence |
|---|---|---|
| Endpoint PCR genotyping | Exon 16 forward | ACGTTCAAGCTACCAGTGGG |
| | Exon 16 reverse | GGTTTCCTGGTGCTGTCAGT |
| | CARM1 WT | CTGCCATCTCTATGGCCTCG |
| | CARM1 SA | GGGATGGACATGGGGGCA |
| qPCR genotyping | qPCR CARM1 forward | CACACGTTCAAGCTACCAGT |
| | qPCR CARM1 reverse | AGGCACCTAACTCCCATAGT |
| | CARM1 WT probe | CATCTCTATGGCCTCGCCCATGTC |
| | CARM1 SA probe | TGCCATCTCTATGGCTGCCC |

**Table S2.** Antibodies used and applications.

| Application | Antibody | Dilution |
|---|---|---|
| Primary antibody (Western blot) | Anti-CARM1 (Bethyl, A300-421A) | 1:2000 |
| | Anti-NFIBme2a (generous gift from Mark Bedford (MD Anderson)) | 1:1000 |
| | Anti-PAX7 (DSHB) | 1:500 |
| | Anti-MYF5 (Santa Cruz, sc-518056) | 1:500 |
| | Anti-MYHC (Sigma, M4276) | 1:1000 |
| Secondary antibody (Western blot) | Goat Anti-Rabbit IgG (H + L)-HRP Conjugate (Bio Rad, 1706515) | 1:10,000 |
| | Goat Anti-Mouse IgG (H + L)-HRP Conjugate (Bio Rad, 1706516) | 1:10,000 |
| Primary antibody (IF) | Anti-PAX7 (DSHB) | Undiluted |
| | Anti-MYOD (Santa Cruz, sc-377460) | 1:200 |
| | Anti-MYOG (Abcam, ab124800) | 1:200 |
| | Anti-MYHC (Sigma, M4276) | 1:150 |
| | Anti-Laminin (Sigma, L9393) | 1:1000 |
| | Anti-CARM1 (Bethyl, IHC-00045) | 1:200 |
| Secondary antibody (IF) | Donkey anti-Rabbit IgG (H+L), Alexa 647, (Invitrogen, A32795) | 1:1,000 |
| | Donkey anti-rabbit IgG (H+L) Alexa 488 (Invitrogen, A21206) | 1:1000 |
| | Donkey anti-Mouse IgG (H+L), Alexa 647, (Invitrogen, A3278) | 1:1,000 |
| | Goat anti-mouse IgG2b Alexa 555 (Invitrogen, 421147) | 1:1000 |
| | Goat anti-mouse IgG1 Alexa 647 (Invitrogen, A21240) | 1:1000 |
| Positive selection (FACS) | a7-integrin-Alexa 647 (UBC Ablab, 67-0010-05) | 1:1,000 |
| | VCAM-PE/Cy7 (BioLegend, 105720) | 1:400 |
| Negative selection (FACS) | CD31-PE (BD Biosciences, 553373) | 1:10,000 |
| | CD45-PE (BD Biosciences, 553081) | 1:10,000 |
| | CD11b-PE (BD Biosciences, 553311) | 1:10,000 |
| | Ly6A/E (Sca-1)-PE (BD Biosciences, 553108) | 1:10,000 |

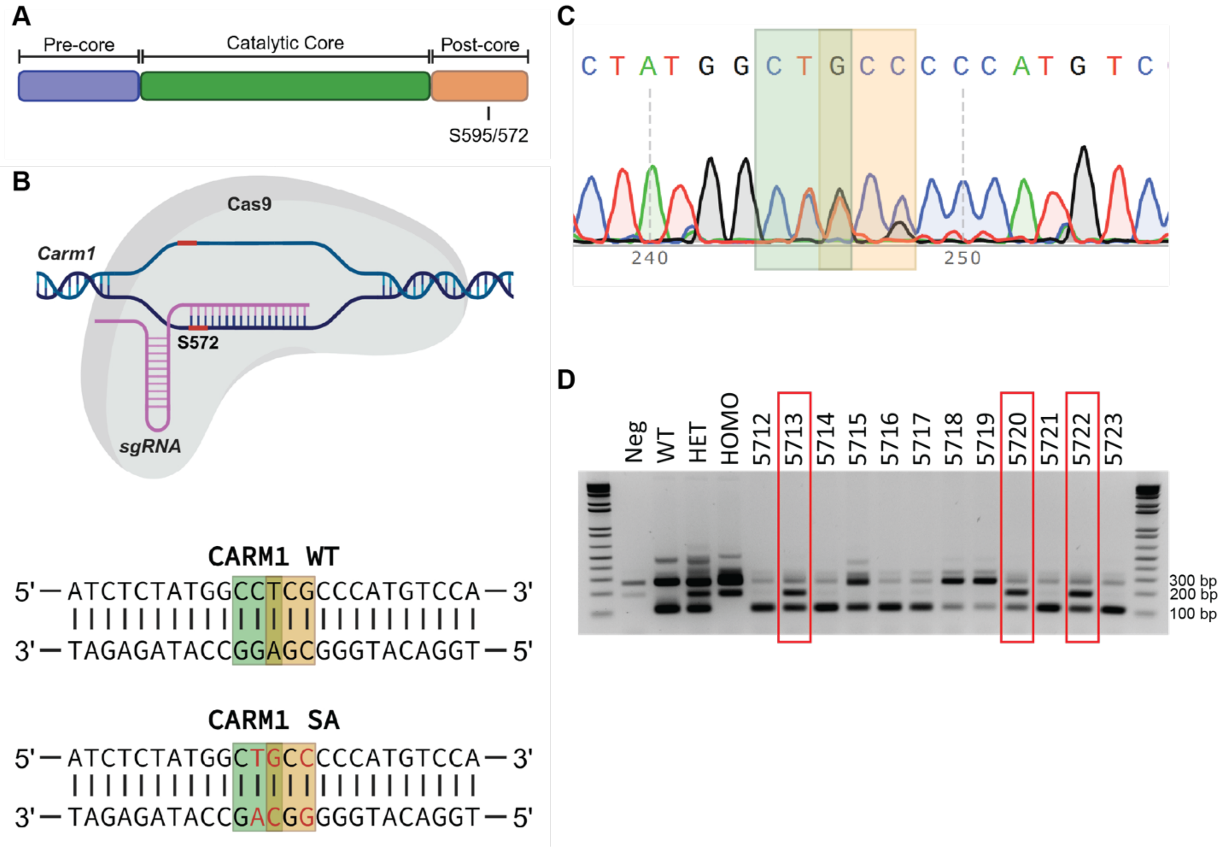


**Figure S1. CRISPR/Cas9-mediated mutagenesis of the CARM1 Ser572/595 residue. A)** Schematic representation of the CARM1 protein indicating the location of Ser572/595, the residue phosphorylated by p38γ/MAPK12. **B)** Nucleotide sequence of *Carm1* surrounding the Ser572/595-encoding codon. CRISPR/Cas9-mediated genome editing was used to introduce a serine-to-alanine substitution at this site. Mutated nucleotides are highlighted in red, the PAM sequence is indicated by a green box, and the Ser572/595 codon is indicated by an orange box. **C)** Representative Sanger sequencing chromatograms from heterozygous CARM1 SA mice confirming the mutation. The PAM sequence is indicated by a green box and the Ser572 codon by an orange box. **D)** Endpoint PCR genotyping assay used to distinguish CARM1 WT and CARM1 SA mice. The WT allele produces a 100 bp fragment, whereas the CARM1 SA allele produces a 180 bp fragment. The Exon 16 amplicon is 300 bp. Successful CARM1 SA heterozygous mice are highlighted in red.

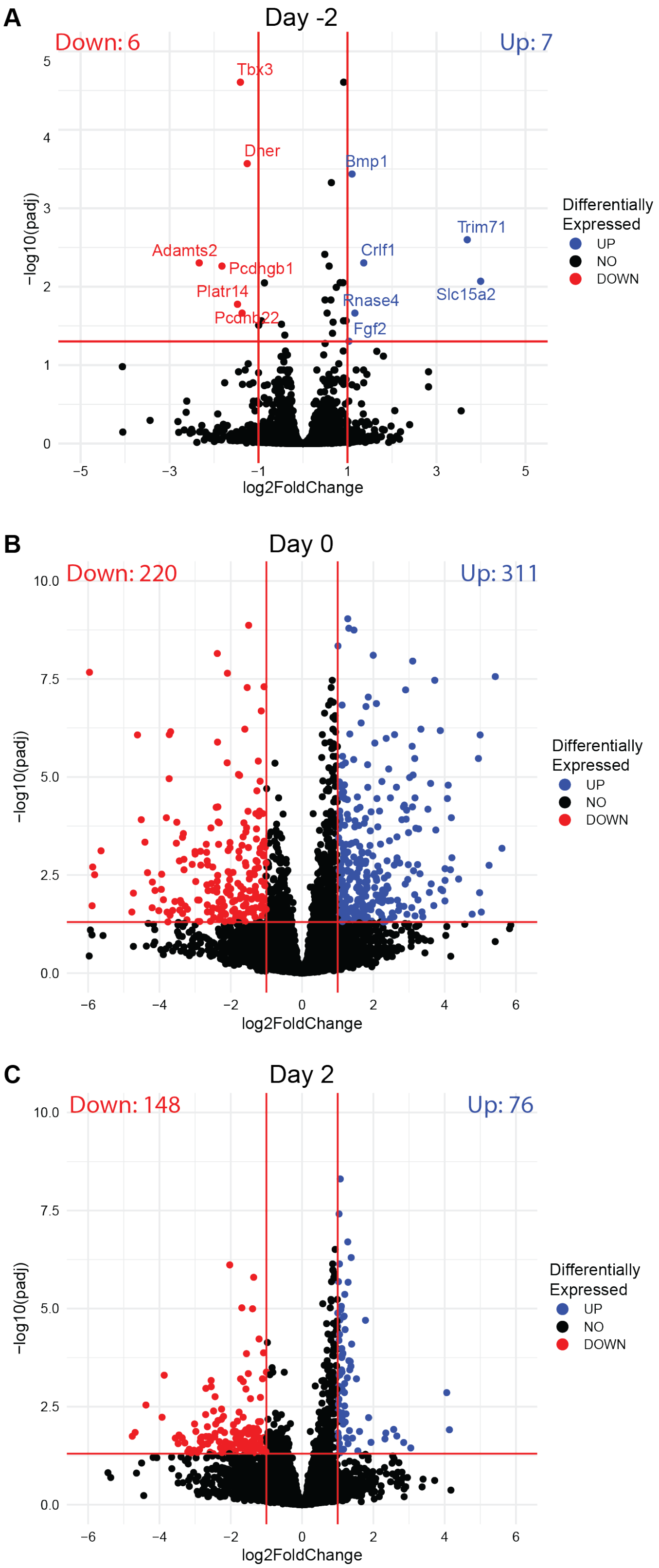


**Figure S2. Transcriptomic profiling of CARM1 WT and SA myoblasts during myogenic differentiation.** Volcano plots showing differentially expressed genes (DEGs; adjusted $p < 0.05$) in CARM1 SA relative to WT myoblasts at **A)** basal (day -2), **B)** day 0, and **C)** day 2 of differentiation. n = 4 WT and n = 3 CARM1 SA biological replicates.

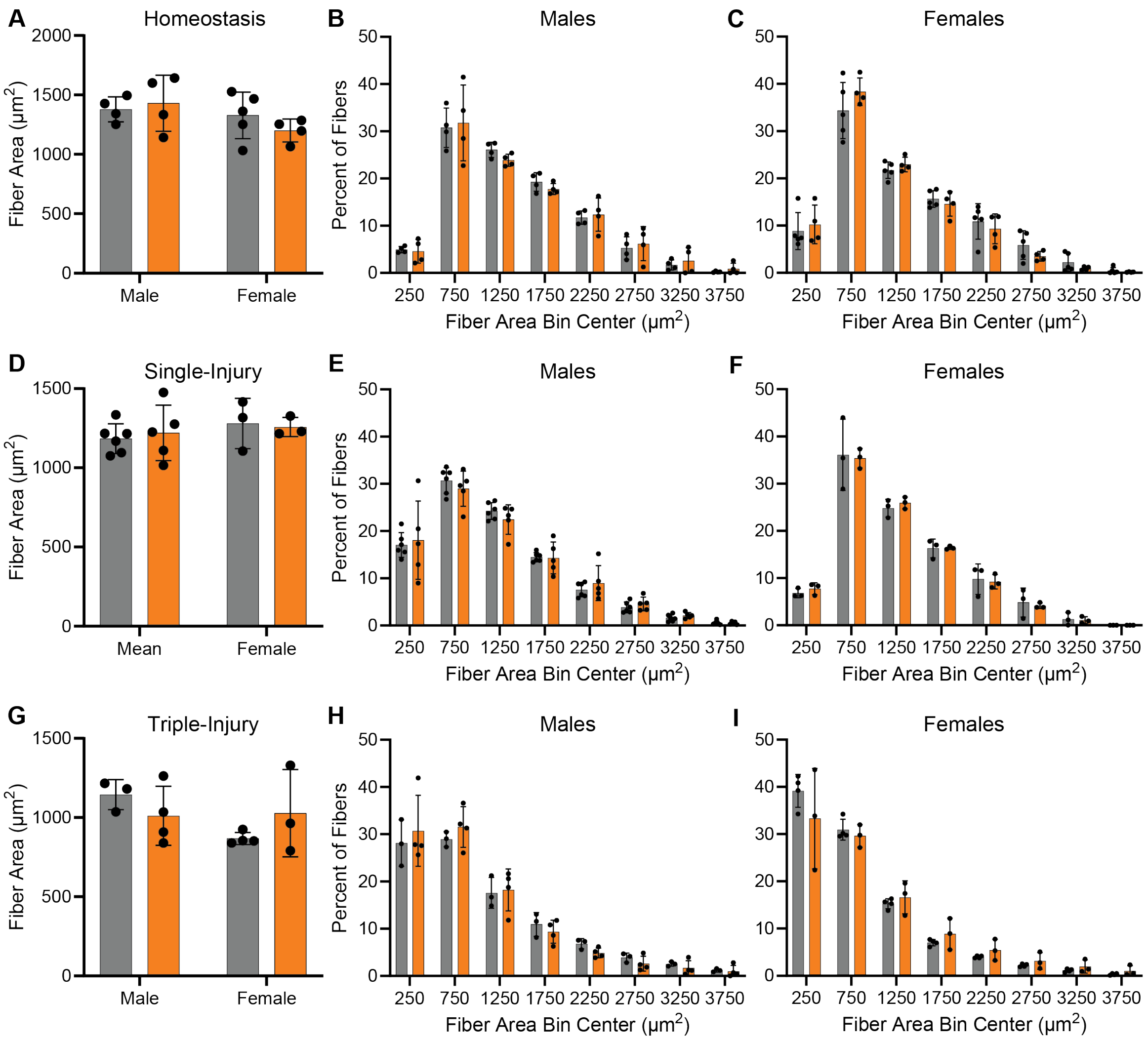


**Figure S3. Muscle fiber cross-sectional area and size distribution in CARM1 wild-type and SA mice.** Mean muscle fiber cross-sectional area was quantified from TA muscle cross-sections of CARM1 WT and SA mice under **A)** homeostatic, **D)** single-injury, and **G)** triple-injury conditions. Frequency distributions of muscle fiber sizes are shown for **B-C)** homeostatic, **E-F)** single-injury, and **H-I)** triple-injury muscles with male (**B**, **E**, **H**) and female (**C**, **F**, **I**) mice displayed separately. Data are presented as mean ± SD, n = 3-6 biological replicates per sex and genotype as indicated; two-way ANOVA with Fisher's LSD post hoc test.

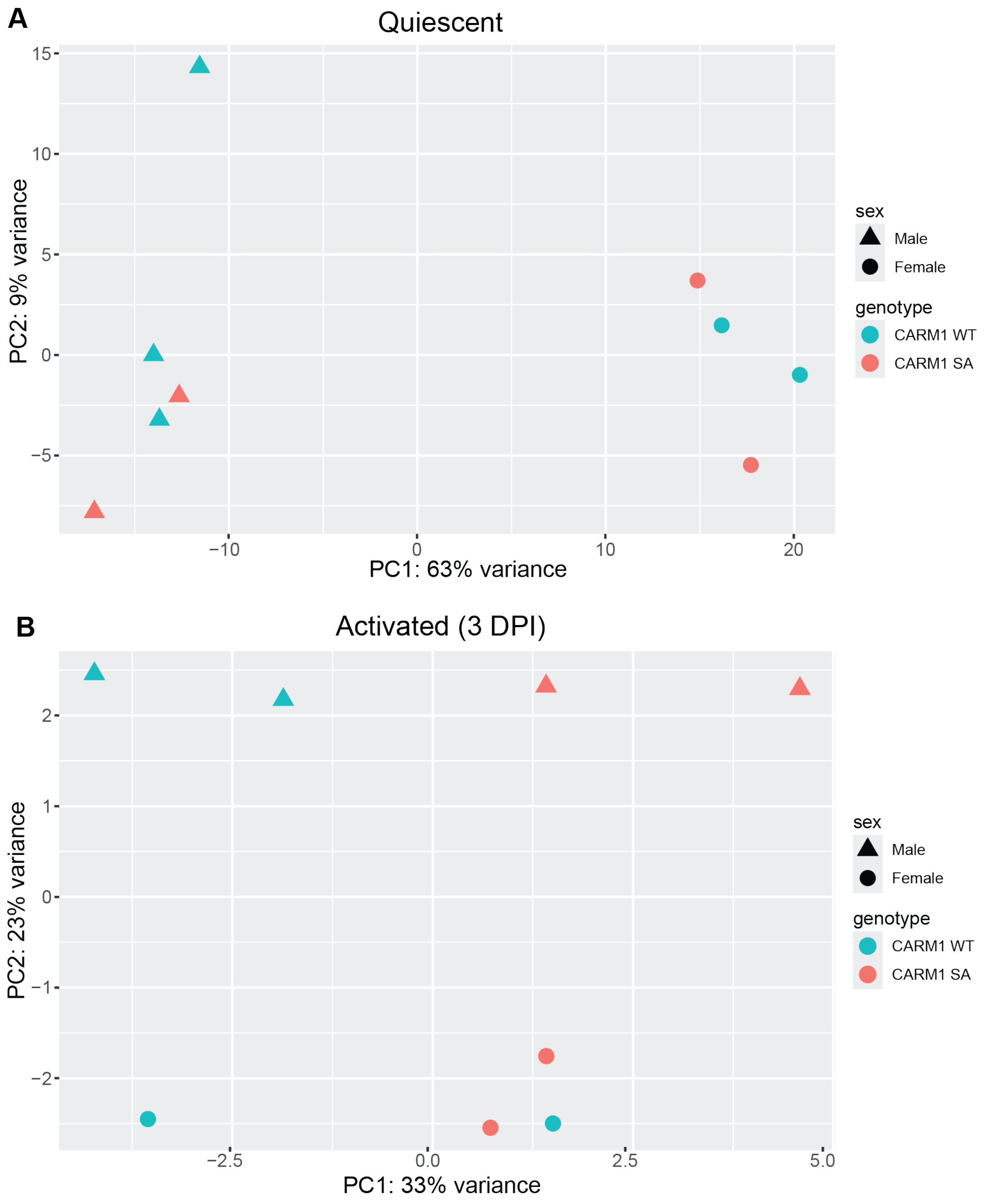


**Figure S4. Principal component analysis of transcriptomic profiles from quiescent and activated satellite cells in CARM1 wild-type and SA mice.** Principal component analysis (PCA) of bulk RNA sequencing data from **A)** quiescent satellite cells isolated from homeostatic muscle and **B)** activated satellite cells isolated from regenerating muscle at three days post-injury (3 DPI). Samples are colored by genotype and shaped according to sex. For **A**, n = 5 CARM1 WT (3 male, 2 female) and n = 4 CARM1 SA (2 male, 2 female). For **B**, n = 4 for each genotype (2 male, 2 female).
